# Physiological and behavioural resistance of malaria vectors in rural West-Africa : a data mining study to address their fine-scale spatiotemporal heterogeneity, drivers, and predictability

**DOI:** 10.1101/2022.08.20.504631

**Authors:** Paul Taconet, Dieudonne Diloma Soma, Barnabas Zogo, Karine Mouline, Frederic Simard, Alphonsine Amanan Koffi, Roch Kounbobr Dabire, Cedric Pennetier, Nicolas Moiroux

## Abstract

Insecticide resistance and behavioural adaptation of malaria mosquitoes affect the efficacy of long-lasting insecticide nets - currently the main tool for malaria vector control. To develop and deploy complementary, efficient and cost-effective control interventions, a good understanding of the drivers of these physiological and behavioural traits is needed. In this data-mining exercise, we modelled a set of indicators of physiological resistance to insecticide (prevalence of three target-site mutations) and behavioural resistance phenotypes (early- and late-biting, exophagy) of anopheles mosquitoes in two rural areas of West-Africa, located in Burkina Faso and Cote d*’*Ivoire. To this aim, we used mosquito field collections along with heterogeneous, multi-source and multi-scale environmental data. The objectives were i) to assess the small-scale spatial and temporal heterogeneity of physiological resistance to insecticide and behavioural resistance phenotypes, ii) to better understand their drivers, and iii) to assess their spatio-temporal predictability, at scales that are consistent with operational action. The explanatory variables covered a wide range of potential environmental determinants of vector resistance to insecticide or behavioural resistance phenotypes : vector control, human availability and nocturnal behaviour, macro and micro-climatic conditions, landscape, etc. The resulting models revealed many statistically significant associations, although their predictive powers were overall weak. We interpreted and discussed these associations in light of several topics of interest, such as : respective contribution of public health and agriculture in the selection of physiological resistances, biological costs associated with physiological resistances, biological mechanisms underlying biting behaviour, and impact of micro-climatic conditions on the time or place of biting. To our knowledge, our work is the first modeling insecticide resistance and feeding behaviour of malaria vectors at such fine spatial scale with such a large dataset of both mosquito and environmental data.

## Introduction

Malaria remains a major public health concern in Africa, with 234 million cases and 593 000 death over the continent in 2021 (WHO, 2022). After years of steady reduction in the disease transmission mainly due to the scale-up of vector control (VC) interventions (in particular insecticide-based tools such as long lasting insecticide nets (LLIN) and indoor residual spraying (IRS)) (Bhatt et al., 2015), progress is now stalling since 2015 (WHO, 2022). Involved in such worrying trends are a combination of biological, environmental and socio-economical factors. The mosquito biology, with the buildup of adaptive changes in the mosquito vectors populations enabling them to avoid or circumvent the lethal effects of insecticides, is most likely playing a very important contribution (Killeen, 2014). These changes are framed as vector *resistance* to insecticides. As a consequence of the widespread use of insecticides (in agriculture and public health), vector resistance has arisen rapidly in malaria vectors in many areas of Africa and above (Durnez and Coosemans, 2013; Riveron et al., 2018); and as previously indicated, is now at such level that it compromises the effectiveness of the most efficient malaria control interventions (Gatton et al., 2013; Hemingway et al., 2016; Killeen, 2014; Sokhna et al., 2013). Complementary and locally-tailored VC strategies taking into account the great diversity of vectors resistance mechanisms (see below) are therefore needed to target these vectors contributing to residual malaria transmission (Corbel and N*’*Guessan, 2013; Durnez and Coosemans, 2013; Hemingway et al., 2016; Moiroux, 2012; Riveron et al., 2018; Sokhna et al., 2013; WHO, 2017).

Vector resistances to insecticide are usually split into two categories : *physiological* and *behavioural* resistance (Lockwood et al., 1984; Sokhna et al., 2013). Physiological resistance refers to biochemical and morphological mechanisms (e.g. target-site modifications, metabolic resistance, cuticular thickness) that enable the mosquito to withstand the effects of insecticide despite its contact with it (Davidson, 1957). Among the physiological resistances, the target-site mutations L1014F (*kdr-w*) (Martinez-Torres et al., 1998), L1014S (*kdr-e*) (Ranson et al., 2000), and G119S (*ace-1*) (Weill et al., 2004), conferring insecticide resistance to pyrethroids (*kdr-w* and *kdr-e*) and to carbamates and organophosphates (*ace-1*), have been extensively described. behavioural resistance, on its side, refers to any modification of mosquito behaviour that facilitates avoidance or circumvention of insecticides (Carrasco et al., 2019; Gatton et al., 2013; Riveron et al., 2018). behavioural resistance of mosquitoes to insecticides can be qualitative (i.e. modifications that prevent or limit the contact with the insecticide) or quantitative (i.e. modifications that stop, limit or reduce insecticide action once contact has occurred, e.g. escaping, behavioural thermoregulation or curative self-medication) (Carrasco et al., 2019). Up-to-date, the behavioural resistance mechanisms described in the literature are mainly qualitative and consist in spatial, temporal, or trophic avoidance. In particular, in the anopheline populations, the following behavioural qualitative resistance mechanisms have been described after the scale-up of insecticide-based VC tools (Durnez and Coosemans, 2013) : i) increase of exophagic or exophilic behaviours (spatial avoidance), where mosquitoes shifted from biting or resting indoor to outdoor, ii) increase of early- or late-biting behaviours (temporal avoidance), where mosquitoes shifted from biting at night to earlier in the evening or later in the morning, iii) increase of zoophagic behaviours (trophic avoidance), where mosquitoes shifted from biting on humans to biting on animals.

To help develop and deploy complementary VC strategies that are efficient and cost-effective, a better understanding of the spatiotemporal distribution and drivers of both vector physiological resistance and feeding behaviour is needed at a local scale. We raise here a set of questions that, among others, must be explored further at local scale towards this aim :

### > What is the respective contribution of public health and agriculture in the selection of physiological resistances in Anopheles vectors ?

The molecular and genetic basis of physiological resistance has been widely acknowledged: under the pressure of insecticides, mutations that enable the vectors to survive are naturally selected and then spread over the generations (Labbé et al., 2017; Martinez-Torres et al., 1998). The main force that governs the selection of a physiological mechanism of resistance in a population of insects is therefore the pressure induced by insecticide exposure. This pressure can be induced by the vector control tools, or by the runoff of pesticides used in agriculture (in many cases, the same as those used for impregnation of bed nets) into the malaria vectors breeding sites (Chandre et al., 1999; Hien et al., 2017; Reid and McKenzie, 2016; Yadouleton et al., 2011). Assessing the relative contribution of these two pressures on the selection of resistant phenotypes is critical to further predict the relative impacts of public health and agriculture on the growth of physiological resistances and act consequently.

### > What are the biological mechanisms underlying behavioural resistances ?

Contrary to physiological resistance, the biological mechanisms underlying behavioural resistance are still poorly known (Carrasco et al., 2019; Durnez and Coosemans, 2013; Killeen, 2014; Main et al., 2016). In particular, a pending question, having important implications for vector control, is whether behavioural shifts reflect evolutionary adaptations in response to selection pressures from vector control tools, as for physiological resistances (*constitutive resistance*) or are manifestations of pre-existing phenotypic plasticity which is triggered when facing the insecticide or in response to environmental variation that reduces human host availability (*inducible resistance*). Inducible resistance imply that vectors rapidly revert to baseline behaviours when VC interventions are lifted, whereas constitutive resistance might progressively and durably erode the effectiveness of current VC tools. Understanding the biological mechanisms underlying behavioural resistances is therefore important to assess the long-term efficacy of insecticide-based VC interventions.

### > Are mosquito biting behaviours modulated by local-scale environmental conditions other than insecticide-related ones ?

As aforementioned, the overall rise of behavioural resistances is likely caused by the widespread of insecticide-based vector control interventions. However, local environmental conditions can modulate vector behaviours at the time of foraging activity. Local climatic conditions – e.g. wind, rain, temperature, humidity, luminosity - may for example affect the timing and location of vector biting, as it has been noted in some studies (Kirby and Lindsay, 2004; Kreppel et al., 2020; Ngowo et al., 2017). Mosquitoes with natural endophagic / endophilic preferences might, for example, bite or rest outside if temperature inside is too high or humidity too low, in order to decrease their risk of desiccation-related mortality (Kreppel et al., 2020; Ngowo et al., 2017). Land cover, as well, can affect biting rhythms. It has been noted for example that distance to breeding sites may influence nocturnal host-seeking behaviour, with vectors biting on average earlier in the night in households located close to the breeding sites (Njan Nloga et al., 1993; Snow and Gilles, 2002). Assessing whether and to which extent behavioural resistance traits are influenced by local environmental (climatic or landscape) settings may help design VC tools exploiting the vulnerabilities of vectors.

### > Are there associations between behavioural and physiological resistances ?

Physiological and behavioural resistances may likely coexist in mosquito populations, with the first possibly influencing the second. In fact, physiologically resistant mosquitoes may, theoretically, use the recognition of insecticide-based control tool as a proxy for host presence (framed as *behavioural exploitation* (Carrasco et al., 2019)). Several studies have actually showed that the kdr mutation can modify the host-seeking or biting behaviour of *Anopheles* in presence of insecticide-treated net (Diop, Moiroux, et al., 2015; Diop, Chandre, et al., 2021; Porciani et al., 2017). Such behavioural exploitation could potentially lead to a better host recognition/localization and have a dramatic impact, with the control intervention having the opposite effect to the one expected. It is hence important to assess if and to which extent physiologically resistant mosquitoes exhibit different biting behaviours than their susceptible counterparts.

### > Which adaptative strategy (physiological or behavioural resistance) arises faster ?

Understanding the relative capacity of mosquitoes to develop physiological resistance and to shift their behaviour in response to vector control is necessary to highlight where and when mitigation efforts should be prioritized (Sanou et al., 2021). After introduction / re-introduction of insecticide-based tools, if vectors rapidly shift their behaviour to feed outside or at times when people are not protected by an LLIN, interventions that target such mosquitoes should be quickly deployed. In contrast, the rapid emergence of physiological resistance in vectors who continue to feed indoors and at night indicates that switching to alternative insecticide classes in indoor-based interventions may have a greater impact. Additionally, for a given environment, assessing the relative rate of selection of physiological and behavioural resistances is of direct epidemiological importance : it has been showed for example that under a scenario where LLIN and IRS are both heavily used, changes in the susceptibility to insecticide is likely to have a bigger epidemiological impact than changes in biting times (Sherrard-Smith et al., 2019).

### > Are resistance rates heterogeneous at small spatiotemporal scales ?

Mosquito presence and abundance has already been found heterogeneous in space and time at fine-scale, calling for locally-tailored (species-, season-, and village-specific) control interventions (Moiroux, Bio-Bangana, et al., 2013; Moiroux, Djènontin, et al., 2014; Taconet, Porciani, et al., 2021). However, little is known about the small-scale spatiotemporal heterogeneity of vector resistance. The potential drivers of the selection or triggering of resistant phenotypes (vector control use, land cover, micro-climate, human behaviour, etc.) are likely to vary at small spatiotemporal scales, and so may, at the end of the line, vector resistance. As for abundances, assessing the level of heterogeneity of resistance rates in space and time is important to assess the spatiotemporal scale at which management of vector resistance should be considered.

### > To what extent can we explain and predict vector resistance and biting behaviour in space and time ?

Assessing the levels of explainability and predictability of vector resistance and biting behaviour is important for both scientific and operational purposes. Towards this aim, generating statistical models linking vector resistances or biting behaviours to their potential drivers and assessing their explanatory and predictive powers can help (Shmueli, 2010; Shmueli and Koppius, 2010). High explanatory or predictive powers in the models might suggest that the conditions driving a vector to resist are well understood, and conversely, low explanatory powers might suggest that resistances are driven by factors either yet undiscovered or not included in the models. Additionally, assessing the predictability of resistances in vector populations in space and time is an important step towards mapping vector resistance at every place (e.g. village) and time (e.g. season) in the area, with such decision-support tools important to deploy the right intervention, at the right place and time (Taconet, Porciani, et al., 2021).

In this study, we used field mosquito collections and environmental data collected simultaneously in two rural areas of West-Africa to bring elements of answer to these questions for our areas. Guided by these questions, our overall objectives were i) to assess the fine-scale prevalence and spatiotemporal heterogeneity of physiological resistances and at-risk biting behaviours of malaria vectors in these areas and ii) to better understand their drivers. To do so, we modeled a set of indicators of physiological resistances and behavioural resistance phenotypes (namely *kdr-w*, *kdr-e*, *ace-1* target-site mutations, exophagy, early-biting, and late-biting) at the individual mosquito level using this fine-grained dataset and advanced statistical methods in an exploratory and holistic-inductive approach. Patterns found in the data were interpreted and discussed in light of the topics aforementioned, of importance for the management of malaria residual transmission. We concluded with a set of recommendations to manage vector resistances in our study areas.

## Methods

### Entomological and environmental data

The data used in this work were collected in the frame of the REACT project (Soma, Zogo, Somé, et al., 2020; Zogo, DD Soma, et al., 2019). In this projet, a total of fifty-five villages, distributed in two West-African rural areas (∼ 50×50 km each) located in the areas of Diébougou (southwestern Burkina Faso (BF)) and Korhogo (northern Ivory Coast (IC)) were selected according to the following criteria: accessibility during the rainy season, 200–500 inhabitants per village, and distance between two villages higher than two kilometers. After an exhaustive census of the population in these villages at the beginning of the project, entomological and human behaviours surveys were regularly conducted during 15 months (1.25 year) in the Diébougou area and 18 months (1.5 year) in the Korhogo area. Vector control interventions were implemented both as part of the project and of the national malaria control programs (see below). Figure 1 shows the study areas and the corresponding timelines for data collection and vector control interventions. The data table available in Moiroux, Pennetier, et al. (2023) lists the villages included in the study: names, geographic coordinates, vector control interventions implemented in each village. Entomological data were collected in the field, and environmental data were collated from specific devices (see below) or created from heterogeneous field and satellite-based sources. Below is a description of the data used in our work.

**Figure 1.**
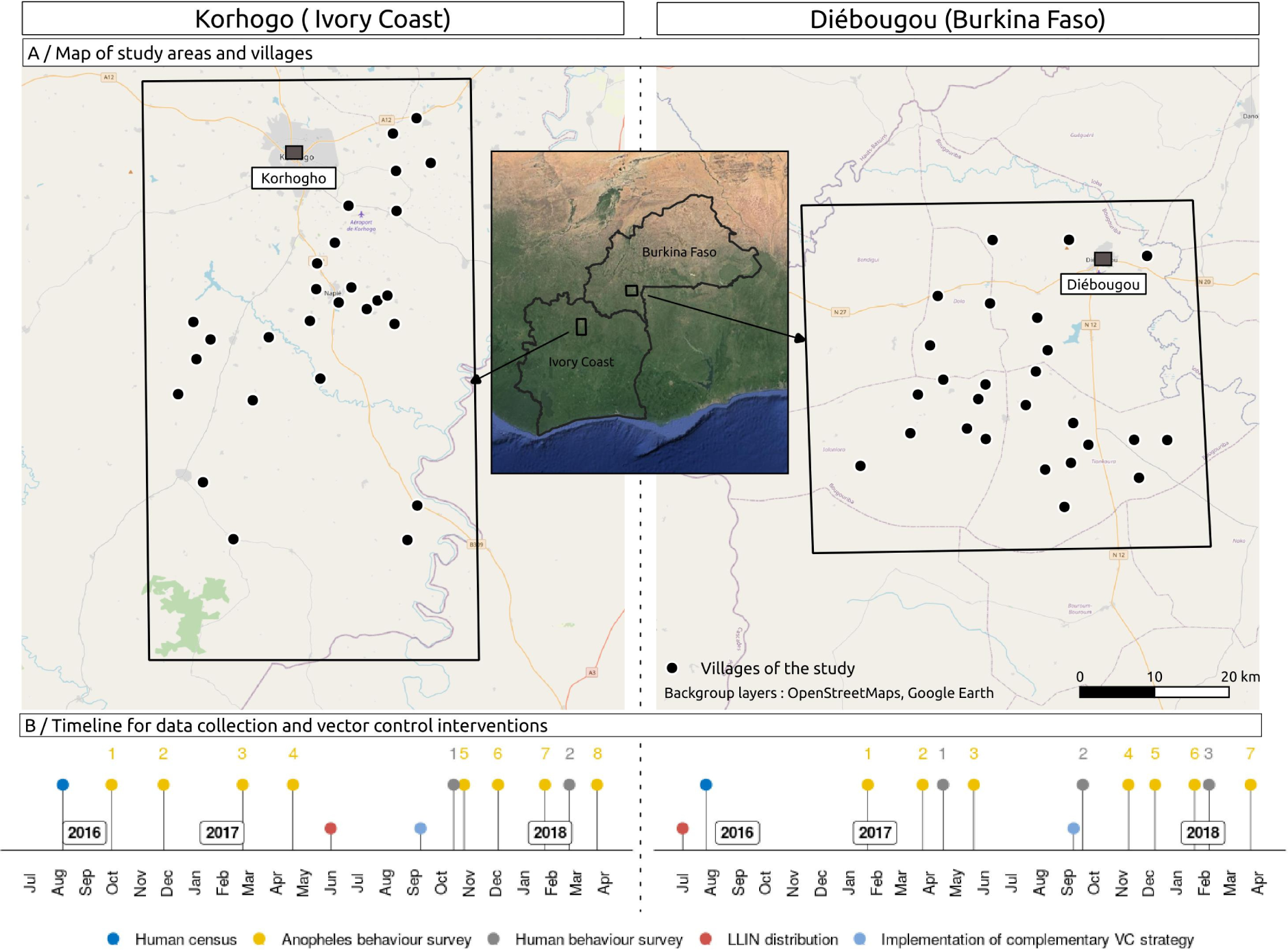
A/ Map showing the study areas and the villages where entomological collections were performed; B/ Timeline for vector control interventions and data collection in the villages. Each color corresponds to a different type of data collected or vector control intervention implemented. The anopheles and human behavioural surveys are numbered.

#### > Anopheles collections

Several rounds of mosquito collections (eight in the Korhogo (IC) area, seven in the Diébougou (BF) area) were conducted in each village. The periods of the surveys span the typical climatic conditions of these tropical areas (except the peak of the rainy season - July to September) (see Additional file 1.A for the spatiotemporal trends of the meteorological conditions). Mosquitoes were collected using the Human Landing Catch (HLC) technique from 17:00 to 09:00 both indoors and outdoors at four sites per village (i.e. eight collection points) for one night during each survey. The distance between indoor and outdoor collection points was at least 10 meters to minimize competition between mosquito collectors (Coffinet et al., 2009). Malaria vectors were identified using morphological keys. All individuals belonging to the *Anopheles funestus* group (in both study areas) and *Anopheles gambiae s.l.* complex (in BF) were identified to the species level using PCR. In IC, due to the very large numbers of *An. gambiae s.l*. vectors collected, a sub-sample only of these individuals (randomly selected in space and time) was identified to species. Finally, in BF, PCR assay were carried out on all the *An. gambiae s.s.* and *An. coluzzii* collected to detect the L1014F (*kdr-w*), the L1014S (*kdr-e*) and the G119S (*ace-1*) target-site mutations. In IC, also due to the large numbers of individuals collected, a subsample only of the *An. gambiae s.l.* were genotyped for the L1014F and G119S mutations. Due to the significant risk of bias associated with the sub-sampling strategy (not all villages were sampled in all surveys), we excluded these data from the analysis. Detailed descriptions of the methods used to obtain these data are provided in Taconet, DD Soma, et al. (2023). These data were published in the Global Biodiversity Information Facility (GBIF) (D Soma et al., 2023).

#### > Data on weather preceding mosquito collections and during mosquito collections

Weather can impact the fitness or the activity of resistant genotypes (Kliot and Ghanim, 2012), as well as the biting behaviour of the mosquitoes (see Introduction). In this work, we recorded or retrieved weather conditions : (i) during mosquito collections (i.e. the HLC sessions), (ii) during the day of collection, and (iii) during the month preceding collection. Weather on the day of collection and during mosquito collection may impact the relative activity of each genotype and phenotypes associated with resistances. Weather during the month preceding the survey, on its side, can impact development and survival rates of both the current and parental generations of collected mosquitoes (Carnevale et al., 2009; Holstein, 1952; Townson, 1993). Regarding our outputs (prevalence of behavioural phenotypes and target-site mutations - see next section), weather during the month preceding collection may therefore impact the fitness of the studied genotypes (for target-site mutations) or possible – and unknown - genotypes associated with studied behavioural phenotypes.

Micro-climatic conditions (temperature, relative humidity, luminosity and atmospheric pressure) were simultaneously recorded where mosquito collections were being conducted. Instruments used to record these data were : for temperature and relative humidity : Hygro Buttons 23 Data Loggers [Proges Plus DAL0084] (temporal resolution (TR): 15 minutes) ; for luminosity : HOBO Pendant® Temperature/Light 8K Data Logger (TR: 15 minutes) ; for atmospheric pressure : Extech SD700 Data Loggers (TR : 10 minutes). Hygro and Hobo loggers were positioned both inside and outside the houses where mosquito sampling was conducted, close to the sampling positions. The barometer was positioned at the center of the village. These field data were completed with satellite or modeled data available at coarse spatial but high temporal resolutions : rain-fall (spatial resolution (SR) : ∼ 11 km, TR : 30 min, source : Global Precipitation Measurement (GPM) IMERG (GSFC, 2019), wind speed (SR : ∼ 28 km,TR : 1h, source : ERA5 (Hersbach et al., 2020)), apparent magnitude of the Moon (SR : 0.001 degrees, TR : 1 day, source : Institute of celestial mechanics and ephemeris calculations).

Meteorological conditions on the day of collection and over one month preceding collection were extracted from satellite imagery. Namely, rainfall estimates were extracted from the GPM - IMERG daily Final products (Center, 2019). Diurnal and nocturnal temperatures were derived from the Moderate Resolution Imaging Spectroradiometer (MODIS) daily Land Surface Temperature (LST) Terra and Aqua products (Wan et al., 2015a,b). These data were then cropped and averaged in 2-km buffer zones around each HLC collection point. From this, variables representing meteorological conditions on the day of collection and over one month preceding collection were constructed (for the latter, by averaging the 30-day time series). Detailed descriptions of the methods used to collect and process these data are provided in Taconet, Porciani, et al. (2021).

#### > Data on host availability and human behaviour

The nocturnal behaviour of humans (hours inside the dwellings, hours of use of LLINs) drives host availability for the mosquitoes and can therefore impact their behaviour. For instance, high LLIN use rate can drive mosquitoes to feed outside, at times when people are not protected, or on alternative sources of blood (Durnez and Coosemans, 2013). Here, human population was counted in each village, through an exhaustive census conducted at the beginning of the project. Then, several human behavioural surveys (two in IC, three in BF) were carried out in each village (see Figure 1). For each survey and village, several households (mean = 14, SD = 2) were randomly selected, and for each household, one to three persons in each age class (0–5 years old, 6–17 years old and ≥ 18 years old) were selected. The head of the household was then asked, for each selected person, on the night preceding the survey : i) whether he/she used an LLIN or not, ii) the time at which he/she entered and exited his own house, and iii) the time at which he/she entered and exited his LLIN-protected sleeping space (where appropriate). Households for human behavioural surveys were independently selected from households for entomological surveys. The surveys were conducted after the distribution of the LLINs (see below), and span the typical climatic conditions of the areas. Detailed descriptions of the methods used to collect these data are provided in Soma, Zogo, Taconet, et al. (2021).

#### > Landscape data

Landscape can have an impact on mosquito foraging behaviour (e.g. the distance to breeding sites can impact biting rhythms) or physiological resistance (e.g. through pesticides used in crops) (see Introduction). Digital land cover maps were produced for each study area by carrying out a Geographic Object-Based Image Analysis (Hay and Castilla, 2008) using multisource very high and high resolution satellite-derived products. From these maps, several variables were derived : the percentage of landscape occupied respectively by cotton fields, by rice fields, and by the other crops (mainly leguminous crops, millet, sorghum) in a 2 km buffer size area around each collection point ; and the distance to the nearest stream (as a proxy for the distance to potential breeding sites, as shown in other studies conducted in these areas (Taconet, Porciani, et al., 2021; Zogo, Koffi, et al., 2019)). For cotton, the variable was binarized as presence / absence of cotton cultivated due to the small range of values. In addition, the geographical location of the households was recorded, and used to derive two indices : the degree of clustering of the households in each village, and the distance from each collection point to the edge of the village. The land cover maps along with detailed descriptions of the methods used to generate them are available at Taconet, Koffi Amanan, et al. (2023) and Taconet, Dabiré, et al. (2023). The methods used to compute the statistical variables from these data are detailed in Taconet, Porciani, et al. (2021).

#### > Vector control

Repeated exposure to insecticides used in vector control interventions is undoubtedly one of the most important drivers of the selection of resistance (see Introduction). In both Burkina Faso and Ivory Coast, LLINs have been universally distributed every 3-4 years since 2010 (PNLP, 2014a,b). In BF, a mass distribution of LLINs (PermaNet 2.0) was carried out by the National Malaria Control Program in July 2016 (i.e. 6 months before our first entomological survey). In IC, our team distributed LLINs in the villages of the project in June 2017 (i.e. height months after the first entomological survey and ten months before the last one). Complementary VC tools were implemented in some of the villages in the middle of the project - namely IRS, ivermectin to peri-domestic animals (IVM), intensive Information Education and Communication to the populations (IEC), and larval control (Larv.) as part of a randomized controlled trial aiming at assessing the benefits of new, complementary VC strategies (Soma, Zogo, Somé, et al., 2020; Zogo, DD Soma, et al., 2019) (see Figure 1, and Additional file 1 available online at this URL (along with the other supplementary material) : https://doi.org/10.23708/VJEEMU (Taconet, D Soma, et al., 2023b)).

## Statistical analyses

### Dependent and independant variables

Six indicators of potential vector resistance to insecticides were modelled :

- three indicators of physiological resistance to insecticide : *kdr-w* mutation, *kdr-e* mutation, *ace-1* mutation,
- three indicators of behavioural resistance phenotypes : early biting, late biting, exophagy. Here, it is unknown whether changes in prevalence of studied mosquito behaviours are the result of constitutive resistances (i.e. inherited traits selected by the insecticide pressure) or of inducible resistance (that rely on phenotypic plasticity). The latter does not fit an accepted definition of insecticide resistance that rely on the inheritance property (Zalucki and Furlong, 2017). Therefore in the remainder of this manuscript, we will qualify the three studied phenotypes, possibly constitutive or inducible, as ***‘***behavioural resistance phenotypes***’***.

Exophagy was defined as the probability for a host-seeking mosquito to bite outdoor (as opposed to indoor). Early biting was defined as the probability for a host-seeking mosquito to bite before 50 % of the LLIN users were declared to be under their bednet in the evening, and late biting was defined as the probability for a host-seeking mosquito to bite after 50 % of the LLIN users were declared to be out of their bednet in the morning (based on the closest - in space and time - human behaviour survey). *Kdr-w*, *kdr-e* and *ace-1* mutations were defined as the probabilities for an allele of a host-seeking mosquito to be mutated (as opposed to wild type). The statistical unit was therefore the mosquito for biting behaviour models and the allele for physiological resistance models. Dependent variables were all binary (0 = absence of resistance/mutation, 1 = presence of resistance/mutation) and models outcomes were probabilities for a mosquito (resp. allele) to be resistant (resp. mutated). Each indicator was modeled separately for each main species in each study area, as determinants of resistance might be species- or site-specific (i.e. mosquitoes might respond differently to environmental variations depending on the species and study area, due to potential local chromosomal forms, adaptation, etc.) (Durnez and Coosemans, 2013; Riveron et al., 2018). As three main species were found in BF and two in IC (see Results section), a total of twenty-one dependent variables were built (exophagy : 3 in BF and 2 in IC ; early biting : 3 in BF and 2 in IC ; late biting : 3 in BF and 2 in IC ; *kdr-w* : 2 in BF ; *kdr-e* : 2 in BF ; *ace-1* : 2 in BF). Based on literature (see Introduction) and available data, we then built independent variables representing potential determinants of each of these resistant phenotypes. These variables are provided in Table 1. To build these variables, the source data were possibly aggregated in space or time, at varying resolutions depending on the considered dependent variable. For example, we constructed a binary variable *“*Rainfall during collection*”* (presence/absence of rainfall during the hour of collection) by summing the source data available at a 30-minutes temporal resolution and then applying a threshold (> 0 mm of rainfall = presence, otherwise absence). Replication data are available online at https://doi.org/10.23708/LV8GEW (Taconet, D Soma, et al., 2023a).

**Table 1.** Independant variables built and their inclusion in the statistical models. A cross (*’*x*’*) indicates that the independent variable (in row) was used as an input in the model (in column). The source data are described in the *’*Entomological and environmental data*’* section of the manuscript, and the binomial statistical models are described in the *’*Statistical analyses*’* section. Some dependent or independent variables, mentioned with a *, were available only in the BF study area.

| Independent variables (in bold : 'family' of variables and source data) | Statistical models (dependent variables) |  |  |  |  |  |
| --- | --- | --- | --- | --- | --- | --- |
|  | Behavioural resistance phenotypes † |  |  | Physiological resistance ‡ |  |  |
|  | Exophagy | Early biting | Late biting | Kdr-w* | Kdr-e* | Ace-1* |
| **Vector control** |  |  |  |  |  |  |
| Vector control tool implemented in the village | x | x | x | x | x | x |
| Time since last distribution of LLIN (months) | x | x | x | x | x | x |
| **Host availability (source : human behavioural surveys)** |  |  |  |  |  |  |
| Number of inhabitants in the village | x | x | x | x | x | x |
| % of the population using an LLIN in the village on the season of collection | x | x | x | x | x | x |
| % of the population indoor (i.e. inside their houses) in the village on the hour of collection | x |  |  | x | x | x |
| % of the population under an LLIN in the village on the hour of collection | x |  |  | x | x | x |
| **Vector resistance / behaviour (source : molecular analyses)** |  |  |  |  |  |  |
| Kdr-e mutation status in the collected mosquito* | x | x | x |  |  | x |
| Kdr-w mutation status in the collected mosquito* | x | x | x |  | x | x |
| Ace-1 mutation status in the collected mosquito* |  |  |  |  | x |  |
| Place of collection of the collected mosquito (indoors or outdoors) |  | x | x |  |  |  |
| **Micro-climatic conditions during collection (source : weather data loggers)** |  |  |  |  |  |  |
| Temperature (°C) | x |  | x | x | x | x |
| Humidity (%) | x |  | x | x | x | x |
| Luminosity (Lux) | x |  | x | x | x | x |
| Atmospheric pressure (hPa) | x |  | x | x | x | x |
| Rainfall (presence/absence) | x |  | x |  |  |  |
| Wind speed outdoor (m/s) | x |  | x |  |  |  |
| Temperature difference between inside and outside the house of collection (°C) | x |  |  |  |  |  |
| Relative humidity difference between inside and outside the house of collection (%) | x |  |  |  |  |  |
| Luminosity difference between inside and outside the house of collection (Lux) | x |  |  |  |  |  |
| Apparent magnitude of the moon on the night of collection (unitless) | x |  |  |  |  |  |
| **Meteorological conditions the day of collection (source : satellite data)** |  |  |  |  |  |  |
| Rainfall on the day of collection (mm) | x | x | x | x | x | x |
| Diurnal temperature on the day of collection (°C) | x | x | x | x | x | x |
| **Meteorological conditions on the month preceding of collection (source : satellite data)** |  |  |  |  |  |  |
| Diurnal temperature (average) on the month preceding collection (°C) | x | x | x | x | x | x |
| Nocturnal temperature (average) on the month preceding collection (°C) | x | x | x | x | x | x |
| Rainfall on the month preceding collection (cumulated mm) | x | x | x | x | x | x |
| **Landscape and crops (source : satellite data)** |  |  |  |  |  |  |
| Degree of clustering of the households in the village) (Clark and Evans aggregation index) | x | x | x |  |  |  |
| Euclidian distance from the collection point to the edge of the village (meters) | x | x | x |  |  |  |
| Euclidian distance from the collection point to the nearest river (meters) | x | x | x |  |  |  |
| % of landscape occupied by rice fields in a 2 km-buffer size area around the collection point |  |  |  | x | x | x |
| Presence / absence of cotton fields in a 2 km-buffer size area around the collection point |  |  |  | x | x | x |
| % of landscape occupied by other types of crops in a 2 km-buffer size area around the collection point |  |  |  | x | x | x |
| **Others** |  |  |  |  |  |  |
| Number of mosquitoes collected at the collection point during the night of collection |  |  |  | x | x | x |

\* BF area only
† Statistical unit = collected mosquito
‡ Statistical unit = allele of collected mosquito

### Modeling workflow

*A graphical representation of the modeling workflow (explained below) is available in Additional* figure 2*. A replication R script (starting from the section ‘Multivariate modeling part 1 : Explanatory model’) is available online at this URL :* https://doi.org/10.23708/LV8GEW (Taconet, D Soma, et al., 2023a).

**Figure 2.**
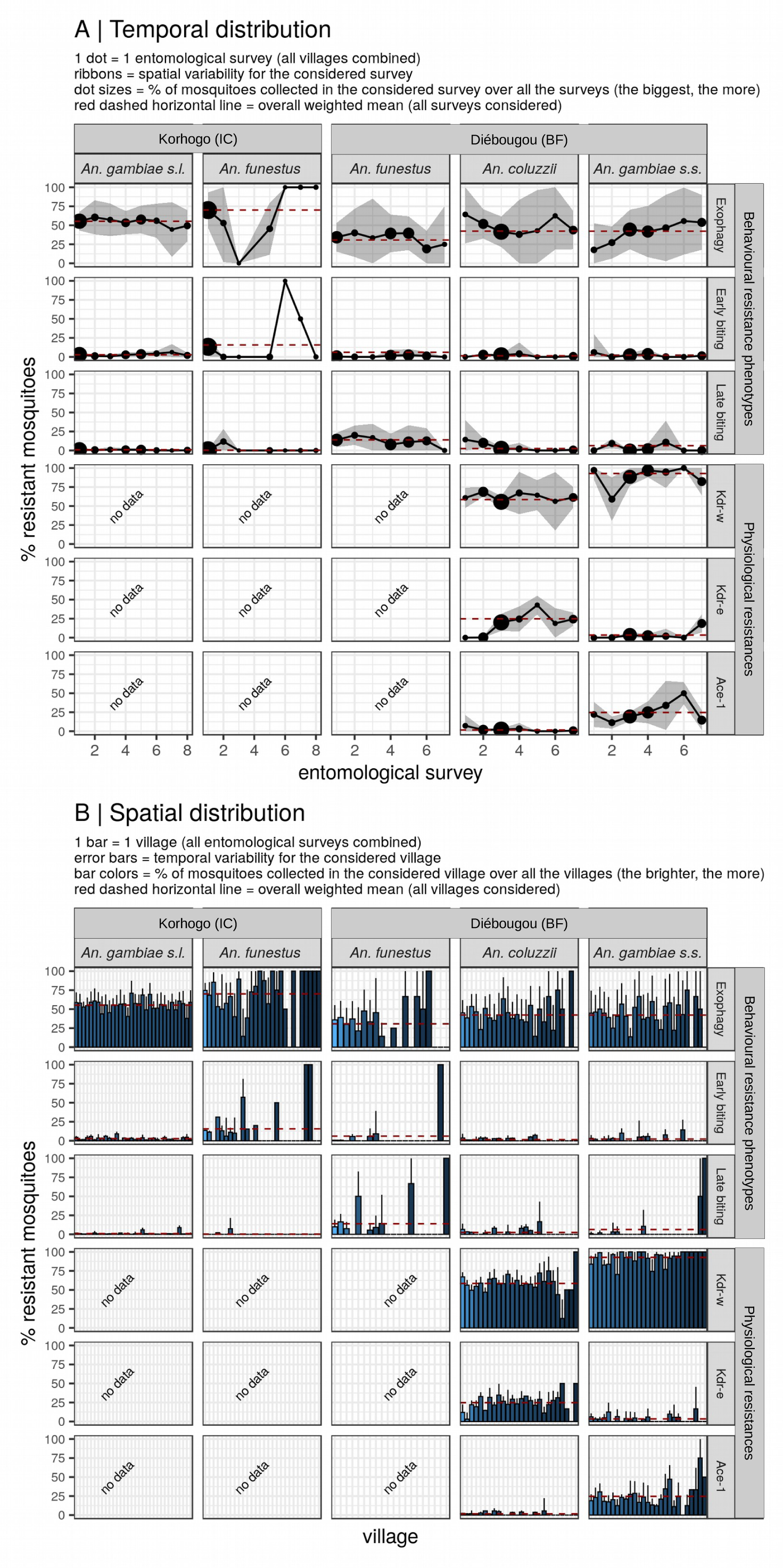
(Previous page) Spatio-temporal distributions of the physiological resistances and be-havioural resistance phenotypes of the main vector species collected (panel A : temporal distribution, panel B : spatial distribution). *For behavioural resistance phenotypes, the y-axis represents the percentage of mosquitoes with resistant phenotypes for the considered survey / village. For physiological resistances, the y-axis represents the allele frequency of the considered mutation for the considered survey / village. Confidence intervals (A : ribbons, B : lineranges) provide indicators of variability of the resistance indicator (A : mean ± standard deviation of the resistance indicator calculated at the village level for the considered entomological survey ; B : mean ± standard deviation of the resistance indicator calculated at the entomological survey level for the considered village). To avoid excessive consideration of small sample sizes, the total number of mosquito collected was represented graphically using the size of dots (A) or the color of the bars (B)*.

#### Pre-processing

First, we excluded from the modeling process those dependent variables that could hardly be modelled due to the combination of very few *‘*resistant*‘* observations and extreme class imbalance (number of samples from the *‘*resistant*‘* class « number of samples from the *‘*sensible*‘* class). The following criteria were used for exclusion: *‘*resistant*’* class *≤* 50 observations & *≤* 3% of the total observations.

Next, we implemented the modeling workflow described below for each remaining dependent variable.

#### Bivariate modeling

We first excluded the independent variables that were poorly associated with the dependent variable (criteria for exclusion : p-value > 0.2 of a bivariate Generalized Linear binomial Mixed-effect Model (GLMM) with nested random effects at the village and collection site level). Next, we calculated the Pearson correlation coefficient among the retained variables and filtered-out collinear variables (correlation coefficient > 0.7) based on empirical knowledge (for instance, diurnal and nocturnal temperature over the month preceding collection were often correlated and in such case we retained nocturnal temperatures ; % of the population indoor and under an LLIN in the village on the hour of collection were often correlated and in such case we retained % of the population under an LLIN). With the set of remaining independent variables, two distinct multivariate models were built, with complementary objectives, as explained in the Box 1 below.

#### Multivariate modeling part 1 : Explanatory model

A binomial GLMM was fitted to the data. Nested random effects were introduced in the model at the village and collection place level. Variables were deleted recursively using an automatic backward variable selection procedure based on the reduction of the Akaike Information Criterion (AIC). Variables belonging to the *‘*vector control*’* (for all resistance models) and *‘*crops*’* (for physiological resistance models only) groups were forced in the multivariate models (i.e. they were not filtered-out in the variable selection procedure) because there are strong *a priori* assumptions associated with these variables. Such variable selection procedure therefore retained all the *‘*vector control*’* and *‘*crops*’* variables (whether significantly associated or not with the dependent variable), and the additional variables that decreased the AIC of the multivariate model.

#### Multivariate modeling part 2 : Predictive model

We additionally fitted a Random Forest (RF) model (Breiman, 2001a) to the data. The model hyperparameters were optimized using a random 5-combinations grid search (Chicco, 2017). Whenever the dependent variable was imbalanced (more than 1/3 imbalance ratio between the positive and negative class), data were up-sampled within the model resampling procedure to cope with well-known problems of machine-learning (ML) models regarding class imbalance (Tyagi and Mittal, 2020).

#### Assessment of effect sizes and significance of independent variables

To interpret the effect of each independent variable in the GLMM model, we plotted, for each independent variable retained in the final model, the predicted probabilities of resistance across available values of that variable (all other things being equals) (i.e. *“*partial dependence plot*”* (PDP) (Friedman and Popescu, 2008)). For reporting and discussion in the manuscript, we kept only variables that had a p-value < 0.05 (results containing the *‘*full*’* models are provided in supplementary material, see Results section). To uncover the possible complex relationships that the RF model had learned, we generated smoothed versions of PDPs for each independent variable. However, we restricted the generation of PDPs to the following cases : i) the Area Under the Receiver Operating Characteristics (AUC) (see below) of the model was > 0.6 (because model interpretation tools of ML models (e.g. PDPs) should be trusted only if the predictive power of the underlying model is good enough (Zhao and Hastie, 2021)) and ii) the range of predicted probabilities of resistance was > 0.05 (i.e. the independent variable, over its range of available values, changed the probability of resistance by at least 5 percentage points).

#### Assessment of models performance

The explanatory power of the GLMM was assessed by calculating the marginal coefficient of determination (R^2^) (Nakagawa and Schielzeth, 2013) from the observed and in-sample predicted values. Marginal R^2^ is a goodness-of-fit metric that measures the overall variance explained by the fixed effects in the GLMM. R^2^ values were interpreted according to the criteria defined by Cohen (2013) : R^2^ *∈* {0; 0.02{ : very weak ; R^2^ *∈* {0.02; 0.13{ : weak ; R^2^ *∈* {0.13; 0.26{ : moderate ; R^2^ *∈* {0.26; 1} : substantial. The predictive power of the RF model was assessed by leave village - out cross-validation (CV), with the Area under the ROC Curve (AUC) chosen as the performance metric. This CV strategy consisted in recursively leaving-out the observations belonging to one village of collection (i.e. the validation fold), training the model with the observations coming from the other villages (i.e. the training fold), and predicting on the left-out set of observations. We hence measured the ability of the model to predict resistance status (*‘*resistant*’* or *‘*non-resistant*’*) on individual mosquitoes caught on new - unseen villages of collection. AUC values were interpreted according to the following criteria : AUC *∈* {0.5; 0.6{ : very weak ; AUC *∈* {0.6; 0.65{ : weak ; AUC *∈* {0.65; 0.75{ : moderate ; AUC *∈* {0.75; 1} : substantial.

##### Box 1

###### What is the difference between explanatory and predictive models, and how were they used for inference in this study ?

Explanatory and predictive models serve distinct but complementary functions in the production of scientific knowledge. In statistics, explanatory modeling refers to *«the application of statistical models to data for testing causal hypotheses about theoretical constructs.»* (Shmueli, 2010). Explanatory modeling, commonly used for inference in many scientific disciplines such as biology or epidemiology, is useful to test existing theories and to reach to “statistical” conclusions about causal relationships that exist at the theoretical level, e.g. : vector control significantly impacts vector resistance (or not). Explanatory modeling needs transparent and interpretable models, such as linear of logistic regression, to extract statistical information about the associations contained in the data (e.g. effect size and statistical significance) and further discuss them. On its side, predictive modeling is *«the process of applying a statistical model or data mining algorithm aimed at making empirical predictions, and then assessing its predictive power.»* (Shmueli, 2010). Predictive modeling requires models capable of capturing complex patterns in the data, including interactions and non-linear associations, such as *machine learning* models like random forests or support vector machines. Predictive analytics is typically recognised for its usefulness in practical applications, such as predicting the incidence of diseases. However, it can also play a crucial role in scientific knowledge production. For instance, predictive models can help generate new theories by capturing and revealing potentially complex, unanticipated patterns within the data. They can as well be used to evaluate the overall relevance of a theory, through the interpretation of the predictive power of the models (Shmueli and Koppius, 2010). In a “big data” context like that of this study, with large datasets containing numerous observations and variables, predictive analytics is increasingly used to support scientific theory development (Breiman, 2001b; Karpatne et al., 2017; Shmueli and Koppius, 2010).

In our study, we use explanatory modeling with GLMMs to i) test whether vector control significantly increases vector resistance, as could be expected, and ii) infer the potential determinants of vector resistance and their effect size. We use predictive modeling with RFs to i) account for potential unhypothesized, complex associations between independent and dependent variables, and ii) infer the overall contribution of the independent variables to the prevalence of vector resistance, allowing

### Software and libraries used

The softwares used in this work were exclusively free and open source. The R programming language (R Core Team, 2018) and the R-studio environment (RStudio Team, 2020) were used as the main programming tools. The QGIS software (QGIS Development Team, 2021) and the *‘*ggplot2*‘* R package (Wickham, 2016) were used to create respectively the map of the study area and the timeline for data collection. The *‘*glmmTMB*’* (Brooks et al., 2017) package was used for the bivariate modeling. The *‘*buildmer*’* package (Voeten, 2020) was used to fit the GLMM models with stepwise selection in the multivariate modeling. The *‘*caret*’* (Wing et al., 2018) and *‘*ranger*’* (Wright and Ziegler, 2017) packages were used to fit the random forest models in the multivariate modeling. The *‘*MLmetrics*’* (Yan, 2016) and *‘*MuMIn*’* (Bartoń, 2020) packages were used to calculate respectively the AUC of the RFs and the marginal R^2^ of the GLMMs. The *‘*jtools*’* (Long, 2020) and *‘*pdp*’* (Greenwell, 2017) packages were used to generate the partial dependence plots of respectively the GLMMs and the RFs. The *‘*broom.mixed*’* (Bolker and Robinson, 2020) package was used to extract the coefficients / odd ratios, confidence intervals and p-values of the multivariate GLMMs. The *‘*patchwork*’* (Pedersen, 2019) and *‘*gridExtra*’* (Auguie, 2017) packages were used to create various plot compositions. The *‘*tidyverse*’* meta-package (Wickham, 2017) was used throughout the entire analysis. A few additional R packages were used to create, tidy, and transform the data used in this work (see (Taconet, Porciani, et al., 2021)). The LibreOffice suite was used to create the tables and other plot compositions.

## Results

### Spatio-temporal heterogeneity of vector abundance

In the Korhogo area (IC), a total of 1792 human-nights of HLC was conducted. A sum of 57722 vectors belonging to the *Anopheles* genus was collected. The main species/complex found were *An. gambiae s.l*. and *An. funestus* (respectively 56267 (97% of all the *Anopheles* collected) and 714 (1%) individuals collected). Among the 56267 *An. gambiae s.l.* collected, 3922 (7%) were identified to species: 3726 (95% of the individual identified to species) were *An. gambiae s.s.* and 196 (5%) were *An. coluzzii*. Hence, in the rest of this article, we will consider the *An. gambiae s.l.* collected in the Korhogo area as *An. gambiae s.s.* In the Diébougou area (BF), a total of 1512 human-nights of HLC was conducted. A sum of 3056 vectors belonging to the *Anopheles* genus was collected. The main species found were *An. coluzzii*, *An. gambiae s.s.* and *An. funestus* (respectively 1321 (43% of all the *Anopheles* collected), 616 (20%) and 708 (23%) individuals collected). As expected, mosquito abundance was heterogeneous in time and space (except for *An. funestus* in IC, for which the vast majority (93 %) of the individuals was collected in the first entomological survey, and almost half of the individuals (42 %) were collected within one single village) (see additional file 1 and additional figure 3 for maps and charts of the spatiotemporal distribution of vector abundance).

**Figure 3.**
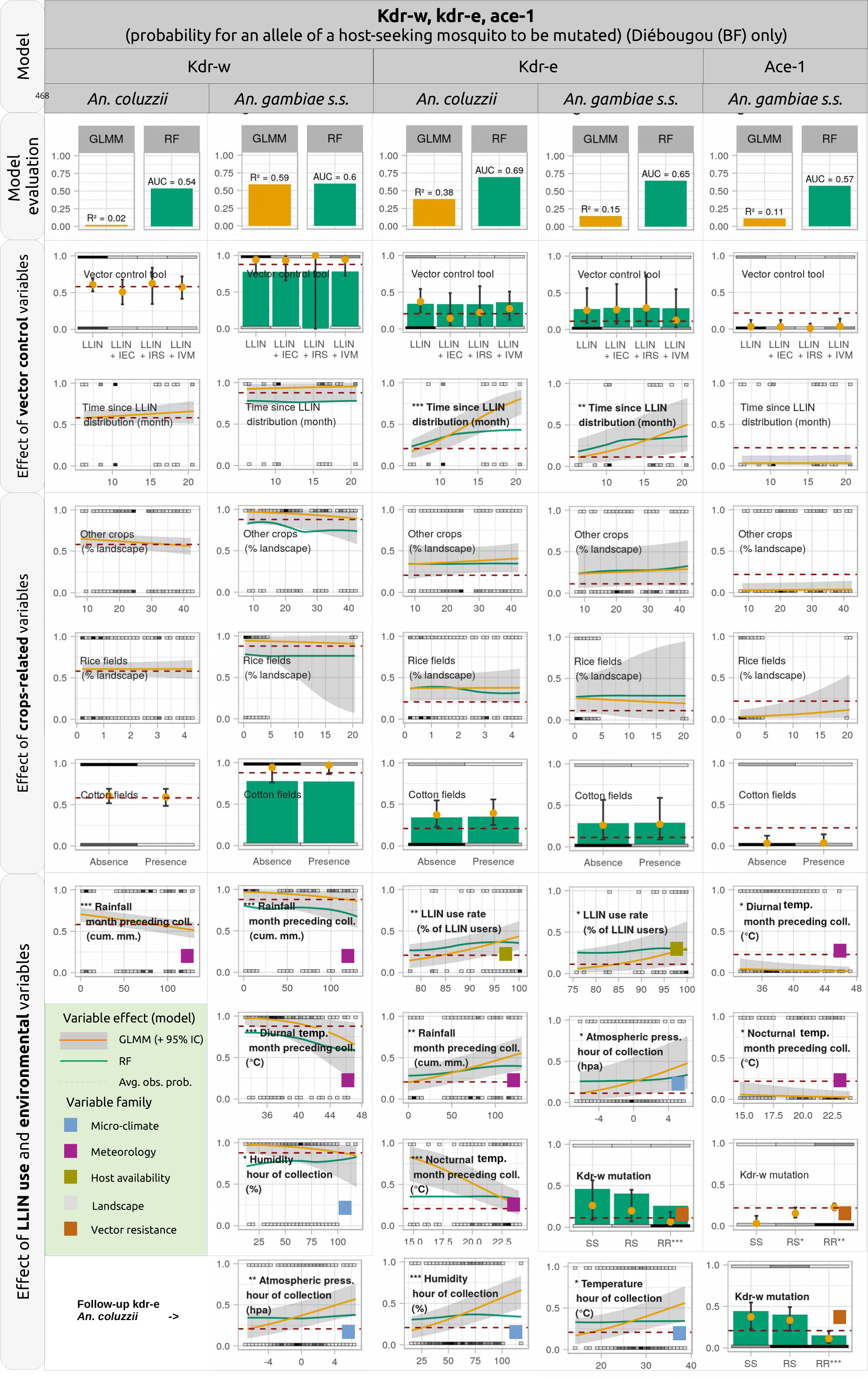
(Previous page) Results of the statistical models of probability of physiological resistance in the malaria vectors. For each model, the top plot shows the explanatory power (R^2^) and predictive power (AUC) of respectively the GLMM and the RF model. The other plots show the predicted probabilities of collecting a resistant vector across available values of each independent variable, holding everything else in the model equal (yellow line: probability predicted by the GLMM model ; green line : probability predicted by the RF model). Non-significant variables (p-value > 0.05) are not presented. Short methodological reminder : vector control and crops variables were forced-in, and the other variables were retained only if they improved the AIC of the model. In addition, for the GLMM models, the other variables were plotted only if their p-value was < 0.05. For the RF models, the predicted probability (i.e. green line) was plotted only if the AUC of the model was > 0.6 and the range of predicted probabilities of resistance for the considered variable was > 0.05. In these plots, the y-axis represents the probability for an allele to be resistant. The red horizontal dashed line represents the overall rate of resistance (see Table 2). The p-values of the GLMMs are indicated through the stars : * : p < 0.05, ** p < 0.01, *** p < 0.001. The coloured squared at the bottom-right represents the ‘family‘ the variable belongs to (one color for each family, see legend inside the light green frame placed on the left hand side of the plot). The grey squares distributed along the x-axis at the top and bottom of each plot represent the measured values available in the data (the darker the square, the more the number of observations) (NB : for atmospheric pressure, the values in the x-axis are centered around the mean)

**Table 2.**
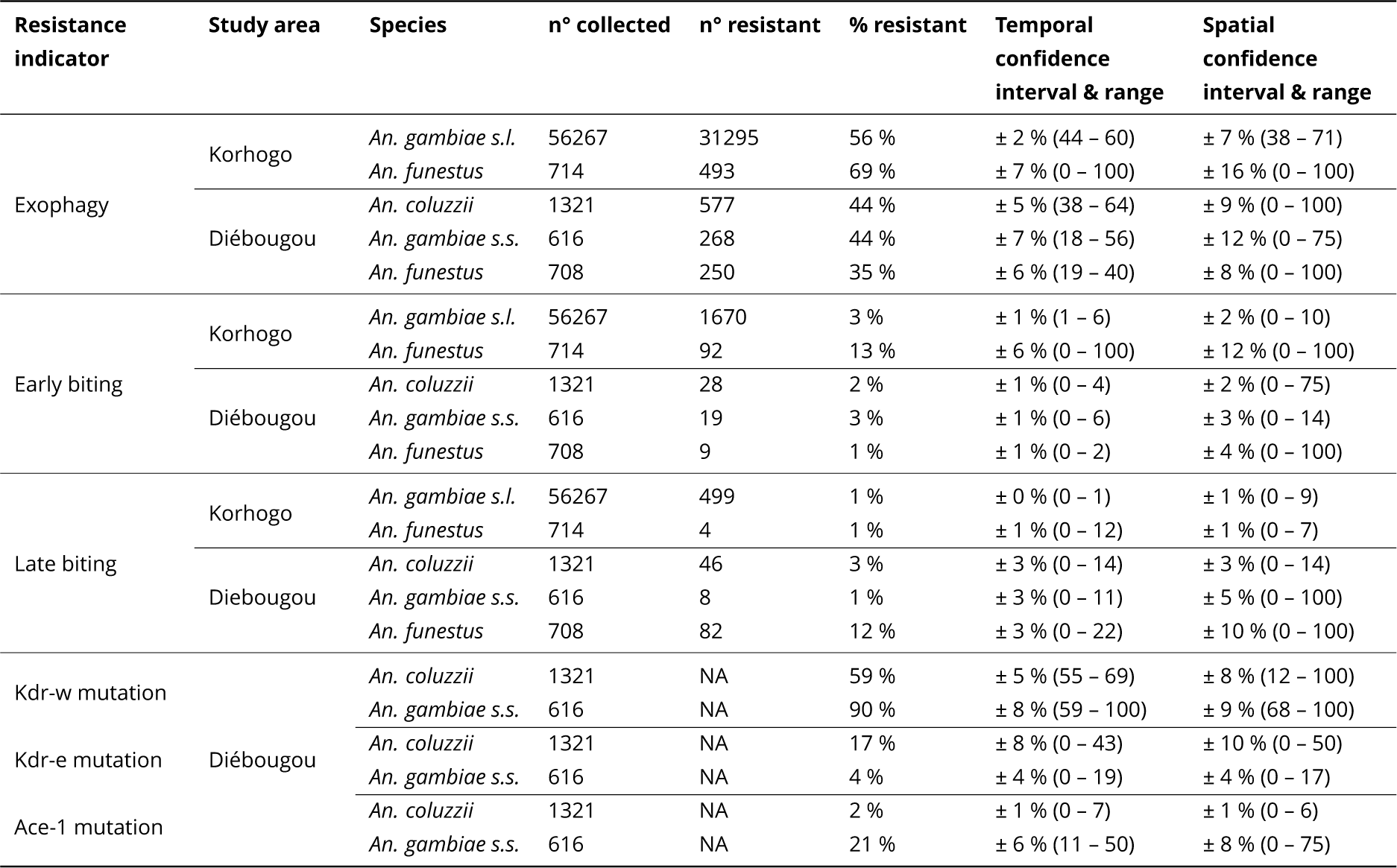
(below) Descriptive statistics for the physiological resistances and behavioural resistance phenotypes of the main vector species collected.

### Spatio-temporal heterogeneity of vector resistance

Table 2 and Figure 2 show, respectively, global and spatiotemporal descriptive statistics on the resistances of the main vector species collected in the two areas.

#### Exophagy rates

In the Korhogo area (IC), the overall exophagy rate (% of bites received outdoor) was 56 % for *An. gambiae s.l.* and 69 % for *An. funestus*. The exophagy rate of *An. gambiae s.l.* varied little, both amongst the entomological surveys and the villages (Temporal Standard Deviation (TSD) (see legend of Table 2 for definition) = ± 2 %, Spatial Standard Deviation (SSD) (see legend of Table 2 for definition) = ± 7 %). The exophagy rate of *An. funestus* was more heterogeneously distributed in time and space (TSD = ± 7 %, SSD = ± 16 %). In the Diebougou area (BF), the overall exophagy rate was 44 % for *An. coluzzii*, 44 % for *An. Gambiae s.s.* and 35 % for *An. funestus*. For the three species, the exophagy rate varied quite sensibly among the entomological surveys (TSD = ± 5%, ± 7%, ± 6% respectively) and the villages (SSD = ± 9%, ± 12%, ± 8% respectively).

#### Early and late biting rates

In the Korhogo area (IC), the early biting rate (i.e. % of bites received before 50% of the LLIN users were declared to be under their bednet at night) was 3% for *An. gambiae s.l.* and 13% for *An. funestus*. The early biting rate was overall stable among the surveys and villages for *An. gambiae s.l.* (TSD = ± 1%, SSD = ± 2%) and was more heterogeneously distributed for *An. funestus* (TSD = ± 6%, SSD = ± 12%). The late biting rate (i.e. % of bites received after 50% of the LLIN users were declared to be out of their bednet in the morning) was lower than the early biting rate : 1% for both *An. gambiae s.l.* and *An. funestus* (only 4 late-bites for *An. funestus*) and was overall stable among the surveys and villages (TSD = ± 0% and SSD = ± 1% for *An. gambiae s.l.*). In the Diébougou area (BF), the early biting rate was respectively 2%, 3% and 1% for *An. coluzzii*, *An. gambiae s.s.* and *An. funestus*. The early biting rate was overall stable among the surveys (TSD = ± 1% for the three species) and to some extent more heterogeneous among the villages (SSD = ± 2%, ± 3%, ± 4% respectively). The late biting rate was respectively 3%, 1% and 12% for *An. coluzzii*, *An. gambiae s.s.* and *An. funestus*. Late biting rates were more heterogeneously distributed than early biting rates, both among the surveys (TSD = ± 3% for the three species) and villages (SSD = ± 3%, ± 5%, ± 10% respectively).

#### Allele frequencies of kdr-e, kdr-w, ace-1 mutations

In the BF area, the allele frequency of the *kdr-w* mutation was 90% for *An. gambiae s.s.* and 59% for *An. coluzzii*. It varied to some extent among the surveys and villages (for *An. gambiae s.s.* : TSD = 8%, SSD = 9% ; for *An. coluzzii* : TSD = 5%, SSD = 8%). The allele frequency of the *kdr-e* mutation was 4% for *An. gambiae s.s.* and 17% for *An. coluzzii*. For *An. gambiae s.s.*, it remained low among the surveys and villages (TSD = SSD = 4%) and for *An. coluzzii*, it varied more sensibly (TSD = 8%, SSD = 10%). The allele frequency of the *ace-1* mutation was 21 % for *An. gambiae s.s.* and 2% for *An. coluzzii*. For *An. gambiae s.s*, it varied sensibly among the surveys and villages (TSD = 6%, SSD = 8%), and for *An. coluzzii* it was overall stably low (TSD = SSD = 1%).

### Dependent variables excluded from the modeling process

Seven of the original twenty-one dependent variables were excluded before statistical modeling due to the very small size of their *‘*resistant*’* class (see Table 2) :

- early-biting in BF for the three species,
- late-biting in BF for *An. coluzzii* and *An. gambiae s.s.*,
- late-biting in IC for *An. funestus*,
- ace-1 in BF for *An. coluzzii*.

### Associations between physiological resistance and environmental variables

For the remaining five models of physiological resistance in the Diébougou area (BF), Figure 3 shows the PDPs of the independent variables retained in the modeling workflow. For the GLMMs, numerical values of odd-ratios, 95% confidence intervals, and p-values are provided in Additional file 4.

#### Associations with variables encoding vector control interventions

No statistically significant association was found between the likelihood of collecting an *Anopheles* carrying any of the target-site mutations and the type of VC intervention (LLIN + complementary tool compared to LLIN only) within the time frame of the study. However, the likelihood of collecting a host-seeking *An. gambiae s.s.* or *An. coluzzii* carrying a resistant *kdr-e* allele increased with the time since LLIN distribution, and as well with the % of users of LLINs in the village population. Noteworthy, for both species the random forest models predicted a significant linear increase in the 12 first months after the distribution, and a slowdown in the increase from the 12th to the 21th month after LLIN distribution. Regarding the others target-site mutations (*kdr-w* or *ace-1*), the likelihood of collecting a host-seeking *Anopheles* carrying them did not increase with the time since LLIN distribution.

#### Associations with variables encoding crops

No statistically significant association was found between the likelihood of collecting a host-seeking *Anopheles* carrying any of the target-site mutations and the % of landscape occupied by crop fields (cotton, rice, or other crops) in a 2 km-wide buffer area around the collection point.

#### Associations with variables encoding micro-climate at the time (hour) of foraging activity

Positive asso-ciations were found between the likelihood of collecting a host-seeking *An. coluzzii* carrying the *kdr-e* mutation and atmospheric pressure, humidity and temperature at the time of collection, as well as that of collecting an *An. gambiae s.s.* carrying the *kdr-e* mutation and atmospheric pressure at the time of collection. A negative association was found between the likelihood of collecting a host-seeking *An. gambiae s.s.* carrying the *kdr-w* mutation and humidity at the time of collection.

#### Associations with variables encoding meteorological conditions during the month preceding collection

Negative associations were found between the likelihood of collecting a host-seeking : *An. coluzzii* carrying the *kdr-w* mutation and cumulated rainfall, *An. gambiae s.s.* carrying the *kdr-w* mutation and both cum. rainfall and mean diurnal temperatures, *An. coluzzii* carrying the *kdr-e* mutation and mean nocturnal temperatures, *An. gambiae s.s.* carrying *ace-1* mutation and both mean diurnal and nocturnal temperatures during the month preceding collection. A positive association was found between the likelihood of collecting a host-seeking *An. coluzzii* carrying the *kdr-e* mutation and cumulated rainfall.

#### Association with variables encoding genotype for other insecticide resistance target-site mutations

The likelihood of collecting a host-seeking *An. gambiae s.s.* or *An. coluzzii* carrying a resistant *kdr-e* allele was negatively associated with the number of mutated *kdr-w* alleles in the collected mosquito. Conversely, the likelihood of collecting a host-seeking *An. gambiae s.s.* carrying a resistant *Ace-1* allele was higher in individuals also carrying *kdr-w* mutated alleles.

### Associations between behavioural resistance phenotypes and environmental vari-ables

For the remaining nine models of behavioural resistance phenotypes, Figure 4 shows the PDPs of the independent variables retained in the modeling workflow. For the GLMMs, numerical values of odd-ratios, 95% confidence intervals and p-values are provided in Additional file 4.

**Figure 4.**
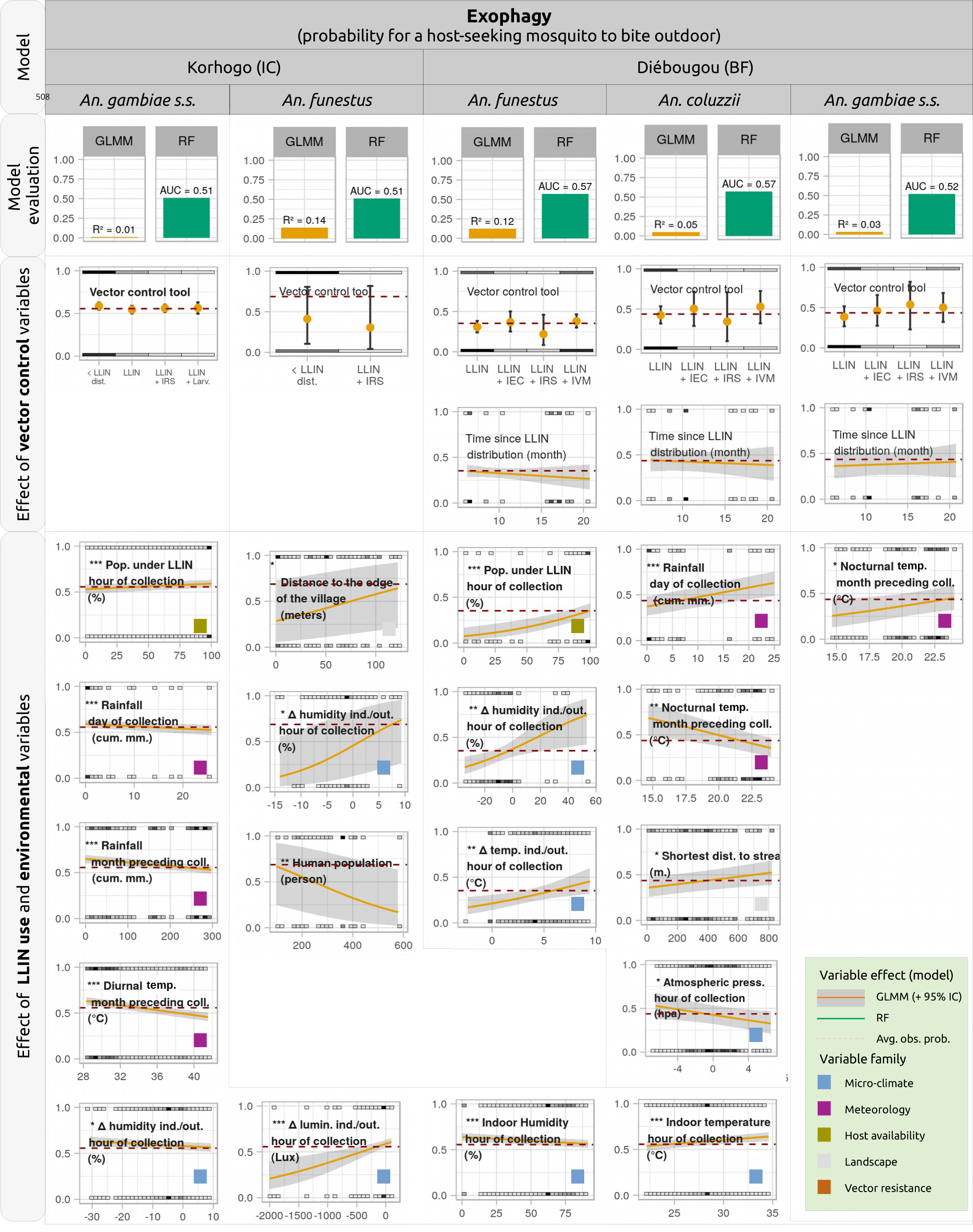
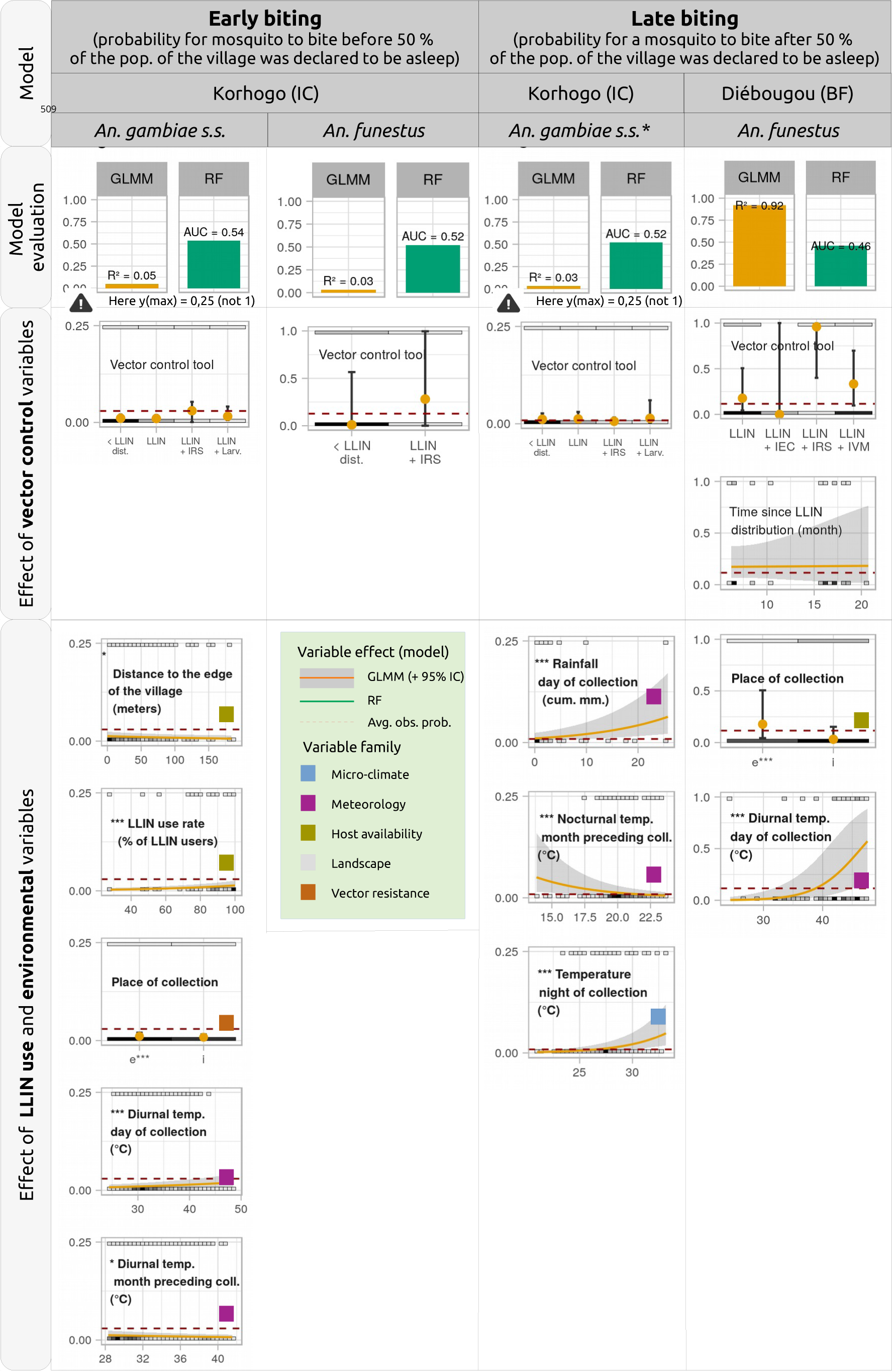
(Previous page) Results of the statistical models of probability of behavioural resistance phenotypes in the malaria vectors. For each model, the top plot shows the explanatory power (R^2^) and predictive power (AUC) of respectively the GLMM and the RF model. The other plots show the predicted probabilities of collecting a resistant vector across available values of each independent variable, holding everything else in the model equal (yellow line : probability predicted by the GLMM model ; green line : probability predicted by the RF model). Non-significant variables (p-value > 0.05) are not presented. Short methodological reminder : vector control variables were forced-in, and the other variables were retained only if they improved the AIC of the model. In addition, other variables were plotted only if their p-value was < 0.05. For the RF models, the predicted probability (i.e. green line) was plotted only if the AUC of the model was > 0.6 and the range of predicted probabilities of resistance for the considered variable was > 0.05. In these plots, the y-axis represents the probability for a mosquito to be resistant. The red horizontal dashed line represents the overall rate of resistance (see Table 2). The p-values of the GLMMs are indicated through the stars * : p < 0.05, ** p < 0.01, *** p < 0.001. The coloured squared at the bottom-right represents the ‘family‘ the variable belongs to (one color for each family, see legend inside the light green frame placed on the left hand side of the plot). The grey squares distributed along the x-axis at the top and bottom of each plot represent the measured values available in the data (the darker the square, the more the number of observations) (NB : for atmospheric pressure, the values in the x-axis are centered around the mean).

#### Associations with variables encoding vector control interventions

No statistically significant association was found between the likelihood of collecting an exophagic, early- or late-biting *Anopheles* and neither the type of VC intervention (LLIN + complementary tool compared to LLIN only) nor the time since LLIN distribution within the time frame of the study.

#### Associations with variables encoding host availability

In the Korhogo area (IC), the likelihood of exophagy of *An. gambiae s.s.* slightly increased with the % of the population under an LLIN at the time of collection. The likelihood of early-biting of *An. gambiae s.s.* increased with the % of users of LLINs in the village population. In the Diébougou (BF) area, the likelihood of exophagy of *An. funestus* increased with the % of the population under an LLIN at the time of collection.

#### Associations with variables encoding landscape

In the Korhogo area (IC), the likelihood of exophagy of *An. funestus* increased with increasing distance to the edge of the village. The likelihood of early-biting of *An. gambiae s.s.* decreased with increasing distance to the edge of the village. In the Diébougou (BF) area, the likelihood of exophagy of *An. coluzzii* increased with increasing distance to the nearest stream.

#### Associations with variables encoding micro-climate at the time (hour) of foraging activity

In the Korhogo area (IC), the likelihood of exophagy of *An. gambiae s.s.* decreased when humidity indoors increased and when humidity got relatively higher indoors compared to outdoors. In addition, it increased when luminosity got relatively higher indoors compared to outdoors. In the Diébougou area (BF), the likelihood of exophagy of *An. funestus* increased when temperature or humidity got relatively higher indoors compared to outdoors.

#### Associations with variables encoding meteorological conditions on the day or night of collection

Positive associations were found between the likelihood of : exophagy of *An. coluzzii* and rainfall (BF area), early-biting of *An. gambiae s.s.* and temperature (IC area), late-biting of *An. gambiae s.s.* and both rainfall and temperature (IC area), late-biting of *An. funestus* and temperature (BF area). A negative association was found between the likelihood of exophagy of *An. gambiae s.s.* and rainfall (IC area).

#### Associations with variables encoding meteorological conditions during the month preceding collection

Negative associations were found between the likelihood of : exophagy of *An. gambiae s.s.* and both cumulated rainfall and mean diurnal temperatures (IC area), exophagy of *An. coluzzii* and mean nocturnal temperatures (BF area), late biting of *An. gambiae s.s.* and mean nocturnal temperature (IC area). A positive association was found between the likelihood of exophagy of *An. gambiae s.s.* and mean nocturnal temperatures (BF area).

#### Associations with variables encoding physiological resistances

As a reminder, the genotypes for the target-site mutations of individual collected mosquitoes were introduced as independent variables in the behavioural resistance phenotypes models in the Diébougou area (BF). Here, these variables were not retained in the variable selection procedure, i.e. no statistically significant association was found between any of the behavioural resistance phenotypes indicators and *kdr-w*, *kdr-e*, or *ace-1* mutations.

### Explanatory and predictive power of the statistical models

Additional figure 5 provides boxplots of observed resistance status vs. predicted probabilities by each model.

#### Exophagy

For the models of exophagy, the explanatory power of the GLMM models was : *‘*very weak*’* for *An. gambiae s.s.* in the Korhogo area (IC), *‘*moderate*’* for *An. funestus* in the Korhogo area (IC)*‘*, weak*’* for *An. funestus*, *An. coluzzii* and *An. gambiae s.s.* in the Diébougou area (BF). The predictive power of the RF models of exophagy was *‘*very weak*’* for all the species in the two study areas.

#### Early and late biting

For the models of early biting, the explanatory power of the GLMM models was *‘*weak*’* for both *An. gambiae s.s.* and *An. funestus* in the Korhogo area (IC). For the models of late biting, the explanatory power of the GLMM was *‘*weak*’* for *An. gambiae s.s.* in the Korhogo area (IC) and *‘*substantial*’* for *An. funestus* in the Diébougou area (BF). The predictive power of the RF models of early and late biting was *‘*very weak*’* for all species in the two study areas, except for the model of late biting of *An. gambiae s.s.* in the Korhogo area (IC) for which it was *‘*weak*’*.

#### Kdr-w, kdr-e, ace-1

For the *kdr-w* mutation in the Diébougou area (BF), the explanatory power of the GLMM models was *‘*weak*’* for *An. coluzzii* and *‘*substantial*’* for *An. gambiae s.s.* ; and the predictive power of the RF models was *‘*weak*’* for *An. coluzzii* and *‘*moderate*’* for *An. gambiae s.s.* For the *kdr-e* mutation in the Diébougou area (BF), the explanatory power of the GLMM models was *‘*substantial*’* for both *An. coluzzii* and *An. gambiae s.s.*; and the predictive power of the RF models was *‘*moderate*’* for *An. coluzzii* and *‘*weak*’* for *An. gambiae s.s.* For the *ace-1* mutation in the Diébougou area (BF), the explanatory power of the GLMM models was *‘*weak*’* for *An. gambiae s.s.* ; and the predictive power of the RF model was *‘*very weak*’*.

## Discussion

In this data mining exercice, we studied indicators of physiological and behavioural resistance phenotypes of several malaria vectors in rural West-Africa at a fine spatial scale (approximately the extent of a health district), using longitudinal data collected in two areas belonging to two different countries, respectively 27 and 28 villages per area, and across 1.25 to 1.5 year. The objectives were to describe the spatial and temporal heterogeneity of vector resistance, and to better understand their drivers, at scales that are consistent with operational action. To our knowledge, our work is the first studying the heterogeneity of vector resistance at such fine spatial scale with such a large dataset of mosquito collection and potential drivers of resistance. In this discussion, we first use our results to provide elements of answers to the questions raised in introduction of this article. We then discuss some implications of the findings for the management of vector resistance in our areas.

### Physiological resistances : potential drivers and spatiotemporal heterogeneity

The main drivers of physiological resistances are insecticides, used either in public health for vector control or in agriculture (see Introduction). In this study, we found that the probability of collecting a host-seeking *An. gambiae s.s.* or *An. coluzzii* in the Diébougou area carrying a *kdr-e* resistant allele significantly increased with both the time since LLIN distribution (up to 12 months after distribution) and the % of LLIN users in the village population. PermaNet 2.0 LLINs have been shown to retain their insecticidal efficacy under field conditions for at least one year after distribution (Djènontin, Alfa, et al., 2023; Kayedi et al., 2017; Kilian et al., 2008; Tan et al., 2016), exerting high selective pressure on vectors over this period at least. In contrast, there was no significant association between any of the target-site mutations and any of the crop-related variable. Altogether, this could indicate that within the spatiotemporal frame of our study, the selection of the kdr-e mutation in the vector population was more likely due to insecticides used in public health than pesticides used in agriculture. In Burkina Faso, pesticides are widely used for cotton and sugar cane (Ouedraogo et al., 2011), but only in lesser proportions in market gardening and cereal production (maize and rice are the only cereals that are treated to a significant extent (MERSI et al., 2016)). Here, in the 2-km wide buffer zones around our collection points crops occupied up to 40 % of the total land, but were mainly made of leguminous crops, millet, sorghum, with cotton and rice being only marginally present. Hence, pesticides are likely not much used (field surveys regarding the use of pesticides by the farmers could confirm this hypothesis). This could explain the absence of association between target-site mutations and the crops-related variables. Noteworthy, the fact that there was no increase in the probability of collecting an *An. gambiae s.l.* carrying a *kdr-e* resistant allele 12 months post-LLIN distribution, as indicated by the RF model, could be attributed to a potential decrease in LLIN insecticidal efficacy after this period (Tan et al., 2016), resulting in lower selection pressure. Finally, we noted that the *kdr-w* and *ace-1* mutations did not increase significantly with the time since LLIN distribution. The absence of increase of the *kdr-w* mutation may be explained by its very high baseline allelic frequencies ; while that of the *ace-1* mutation may be explained by the type of insecticide used to impregnate the LLINs - deltamethrin, which does not select the ace-1 mutation.

The statistical models captured many associations between the likelihood of collecting a physiologically resistant *Anopheles* and the variables encoding weather, both during the month preceding collection and at the hour of collection. These associations could traduce biological costs/advantages associated with target-site mutations, both in terms of fitness and activity, as found elsewhere for other mosquito species (Kliot and Ghanim, 2012). Regarding fitness, we found that the likelihood of collecting a host-seeking mosquito (*An. gambiae s.s.* or *An. coluzzii*) carrying a mutated allele, overall, decreased (to varying extents depending on the species and mutation) when diurnal or noctural temperatures during the month preceding collection got higher, i.e. in the hottest periods of the year (corresponding to ∼ the months of March-April). Carrying a kdr mutation might be associated with a decreased propensity to locate optimal temperatures, potentially resulting in a decreased longevity, fecundity, or ovarian development rates (Foster et al., 2003). Regarding activity, we observed that the likelihood of collecting a mosquito carrying a mutated allele (for the *kdr-e* mutation) decreased when atmospheric pressure, humidity, or temperature at the hour of collection got lower; implying that mosquitoes carrying the *kdr-e* mutation could be less active in colder or drier conditions, or when atmospheric pressure is lower. Noteworthy, our results could also be interpreted in terms of fitness advantages instead of fitness costs: for instance, some studies have highlighted fitness advantages (e.g. for longevity) of the *kdr-w* mutation in *An. gambiae s.l.* in laboratory conditions (Alout et al., 2016; Medjigbodo, Djogbénou, et al., 2021).

We also found interactions between some target-site mutations. Indeed, as the *kdr-e* and *kdr-w* are mutations of the same base pair, the allelic frequency of the *kdr-e* mutation was negatively correlated with the allelic frequency of the *kdr-w* mutation in both *An. gambiae s.s.* and *An. coluzzii*. We also found a positive relationship between the allelic frequencies of the *Ace-1* and *kdr-w* mutations in *An. gambiae s.s.* This is consistent with laboratory observations in *Culex Quinquefasciatus* and *An. gambiae s.s.* showing that the cost of the *Ace-1* mutation is reduced in presence of the kdr mutation (Assogba et al., 2014; Berticat et al., 2008; Medjigbodo, Sonounameto, et al., 2021).

Lastly, we observed that the allelic frequencies of the target-site mutations, within each vector species and for each mutation, were overall quite stable across the villages and seasons within the spatiotemporal frame of the study. At larger spatial and temporal scales, physiological resistances were found more heterogeneous (Moyes et al., 2020). In our study, such homogeneity might be due to a relative homogeneity in space and time of the main determinants of physiological resistance (access and use of insecticide-based vector control interventions). The quite stable rates of physiological resistance throughout the seasons might traduce the fact that the possible fitness costs/advantages are likely rather limited, within the range of meteorological conditions in our area.

### Behavioural resistance phenotypes: potential drivers and spatiotemporal heterogene-ity

An important and pending question is the genetic (constitutive) or plastic (inducible) nature of behavioural resistances (see Introduction). In this study, we found no statistically significant association between any of the indicators of behavioural resistance phenotypes and neither the time since LLIN distribution nor the VC tool implemented. There was hence no evidence of growing frequencies of behavioural resistances (exophagy, early- and late-biting) in response to vector control within the 1.25 to 1.5 years of this study, i.e. no clear indication of constitutive resistance.

Nonetheless, comparison of the exophagic phenotype rates found here with those of previous studies in the same countries, suggests that there may still be a genetic component to mosquito foraging behaviour. Indeed, the exophagy rates measured here tended to be higher than those historically reported for these species. For example, a recent review of *An. gambiae s.l.* biting behaviour from a range of African countries between 2000 and 2018 concluded that during this time period, ∼ 80% of the vectors bite occured indoor (all countries included) and in particular ∼ 75% in Burkina Faso (Sherrard-Smith et al., 2019) (hence respectively ∼ 20% and 25 % outdoor). Here we measured substantially higher levels of exophagy : 44% (range ∼ 18-56%) in the Diébougou (BF) area and 56% (44–60%) in the Korhogo (IC) area. Other recent studies, contemporaneous to ours, have found relatively high levels of exophagy for *An. gambiae s.l.* in rural areas, e.g. 54% in southwestern Burkina Faso (Sanou et al., 2021) or 55% in Ivory Coast (Assouho et al., 2020). Such high levels of outdoor biting, in comparison with past levels, suggest that behavioural adaptations may be ongoing in the study areas, most probably in response to the widespread and prolonged use of insecticide-based vector control tools.

We also found many statistically significant associations between the likelihood of collecting a behaviourally resistant phenotype and the meteorological conditions during the month preceding collection. This might indicate that these phenotypes could be induced by past environmental conditions, acting at the adult or larval stage, or through paternal/maternal effect. Such relationships between environmental condition at the larval stage and adult behaviour have indeed been observed in other insects (Müller et al., 2016, and ref cited in).

The hypothesis of a hereditary component in the behaviour of malaria vectors (at least for the biting hour) is supported by a recent study which has observed, for *Anopheles arabiensis* in Tanzania, that F2 from early-biting F0 (grandmothers) were - to some extent - more likely to bite early than F2 from mid or late-biting F0 (Govella et al., 2023). Under this hypothesis, the relationship between the prevalence of behaviourally resistant phenotypes and the meteorological conditions during the month preceding collection could indicates a cost/advantage, at the adult, larval or both stages, of their associated genotypes.

In our study, the absence of significant association between the probability of behavioural resistances and insecticide-related variables might be due to the relatively short length of the study (2 years). In a similar study conducted in another region of Burkina Faso over a two-year period as well, researchers recorded, as we have, no changes in the biting behaviour of *Anopheles gambiae s.l.*, including early biting, exophagy, and exophily, throughout the duration of the study (Sanou et al., 2021). Although resistance phenotypes can emerge in this time frame (Moiroux, Gomez, et al., 2012), a recent (almost) 4-years-study in Tanzania (Kreppel et al., 2020) detected shifts in vector behaviour (i.e. increased rate of exophily for *An. arabiensis* and *An. funestus*) that could be obscured in shorter-term surveys, in agreement with the hypothesis that mosquito behaviours are likely complex multigenic traits (Main et al., 2016) and might therefore respond slowly to selection (at least, slower than target-site mutations, which are linked to single genes and may hence respond rapidly and efficiently to selection). Anyhow, the results of these various longitudinal studies suggest that long-term monitoring of vector behaviour (> 2 years), particularly in areas with a long history of use of insecticides in public health, is critical to better understand the biological mechanisms underlying behavioural resistances, to potentially assess their development rate, and more broadly to assess residual malaria transmission risk (Durnez and Coosemans, 2013; Kreppel et al., 2020; Sanou et al., 2021).

Weather can impact the fitness of possible genotypes associated with behavioural resistant phenotypes, but may also directly influence the time and location of foraging activity (see Introduction for more details). Here, we found many associations between mosquito host-seeking behaviour and variables representing meteorological conditions on the day or at the hour of collection. For instance, the probability for an *An. gambiae s.s.* to be collected outdoor in the Korhogo area increased when the air indoor was dry, or when the air outdoor became relatively more humid than indoor. Likewise, in the Diébougou area, the probability for an *An. funestus* to be collected outdoor increased when the air outdoor became relatively cooler than indoor. These observations are consistent with the hypothesis of mosquitoes shifting from indoor to outdoor host-seeking in case of desiccation-related mortality risk indoors, as observed and discussed elsewhere (Kessler and Guerin, 2008; Kreppel et al., 2020; Ngowo et al., 2017). The meteorological conditions seemed to cause not only spatial, but also temporal shifts in host-seeking activity. For instance, we found that the probability of collecting a late-biting *An. gambiae s.s.* in the Korhogo area increased when the noctural temperature increased. Several associations also suggest that some malaria vectors may modify their behaviour in response to environmental variation that reduces host availability, as hypothesized elsewhere (Durnez and Coosemans, 2013). For instance, the likelihood of collecting an *An. gambiae s.s.* (in the Korhogo area) or an *An. funestus* (in the BF area) outdoor increased at hours when people were protected by their LLINs. Likewise, the likelihood of collecting an early-biting *An. gambiae s.s.* in the Korhogo area increased when the % of LLIN users in the village increased. Altogether, all these associations suggest that in our study areas mosquito foraging behaviour is driven – to a certain extent - by environmental conditions at the time of foraging activity, i.e. that vectors likely modify their time or place of biting according to climatic conditions or host availability. The many associations that were captured here in field conditions could be further tested experimentally, to quantify their effect more precisely and validate the underlying biological hypothesis.

Although many significant associations between environmental parameters and foraging behaviours have been captured by the models, their explanatory and predictive powers were overall weak. A low explanatory power can indicate either i) that variations in the dependent variable (here, individual vector resistance) are only marginally caused by the independent variables or ii) that the statistical model does not capture properly the true nature of the underlying relationships between the studied effect and its drivers (Karpatne et al., 2017) (e.g. a linear regression cannot, by definition, capture non-linear relationships that might exist in nature). Here, we minimized the risk of omitting important, complex associations by using, complementarily to the binomial regression model, a machine-learning model (namely a random forest) that is inherently able to capture complex patterns contained in the data if any (e.g. non-linear relationships, interactions) (Breiman, 2001a). Still, the models had low predictive powers. Altogether, these results indicate that very likely, despite the amount, granularity and diversity of potential factors measured and introduced in the models, most of the factors driving the individual host-seeking behaviours of the mosquitoes were not introduced in the models. Another possibility could be that some of our independent variables did not represent the actual *“*reality*”* in the field (e.g. the distance to the nearest stream is not necessary an ideal proxy for the distance to the breeding site). Nevertheless, since we used a wide range of variables encoding the environmental conditions at the time of foraging activity, we can hypothesize that within the spatiotemporal frame of the study, mosquito foraging behaviour was only marginally driven by environmental variations. This leaves the floor to other factors, like genetics (see above), learning, or randomness.

To test whether physiological resistance impacts the behaviour of host-seeking mosquitoes, we introduced in the behaviour resistance models of *An. coluzzii* and *An. gambiae s.s.* in the Diébougou area two variables encoding the genotypes for respectively the *kdr-w* and *kdr-e* mutations. No statistically significant association was found. In other words, we could not find, in the field, a behavioural phenotype (among those studied, i.e. exophagy, early- and late-biting) associated with a genotype for one of the target-site mutations. The only study, to our knowledge, having investigated the relationship between the kdr mutation and biting time or location in the field has also reported no statistically significant association between these two mechanisms of resistance to insecticide (Djènontin, Bouraima, et al., 2021). Noteworthy, in our study, there was few variabilities in the genotypes of the collected mosquitoes (i.e. few homozygote susceptible mosquitoes captured, particularly for the *kdr-w* mutation), making it unfavorable to detect assocations between physiological and behavioural resistances. In the Korhogo area, such analysis could not be performed because physiological resistance data was not available at the individual mosquito level.

Finally, we observed that the behavioural resistance phenotypes rates for each vector species in each health district were, overall, relatively homogeneous across the villages and seasons within the spatiotemporal frame of the study (as for physiological resistances). This could mean that the overall dynamics of behavioural resistance occur at broader spatial and temporal scales than those of our study. At larger scales (i.e. among countries and across years in Africa), exophagy rates of *Anopheles* mosquitoes seem, actually, to be more variable (Sherrard-Smith et al., 2019).

### Implications of the findings for the management of vector resistance in the study areas

Long-lasting insecticidal nets have undoubtedly played a major role in reducing malaria cases throughout Africa, thanks both to their barrier and killing effects. More locally, we highlighted the efficacy of their barrier role in the Diébougou area by showing that, for their users, they prevented more than 80% of *Anopheles* bite exposure in the area (Soma, Zogo, Taconet, et al., 2021). However, despite these successes, many studies strongly suggest that the insecticides they are impregnated with are responsible for the rise of physiological resistances in the malaria vectors susceptible populations (Labbé et al., 2017; Riveron et al., 2018) (see Introduction). In our study, the positive and significant associations found between the probability to collect a physiologically resistant mosquito and LLIN-related variables (time since LLIN distribution, LLIN use rate) supports these findings. We also highlighted that in response to an LLIN distribution, physiological resistance seems to grow quite rapidly in a susceptible population. Besides the selection of physiological resistance, comparison with historical data suggests that the vectors may also be progressively changing their feeding behaviour to avoid the effects of the insecticides - although there was no clear evidence of this in the strict context of this study. Such trends in vector resistance may have an important epidemiological impact (Sherrard-Smith et al., 2019). Altogether, these results show, if still necessary, that we urgently need to think more strategically about our use of insecticides in public health tools in our areas. Switching to alternative insecticides, rotating or mixing insecticides, using current or novel insecticides in vector control tools others than long-lasting nets, entirely removing the insecticides from the vector control toolbox, or fostering the use of insecticidal-free tools, are all actions that could be envisaged (Paaijmans and Huijben, 2020). Burkina Faso has, actually, distributed LLINs that mixes pyrethroid with Piperonyl butoxide (PBO) in the last universal LLIN distribution, in 2019.

Here, we observed that both behavioural and physiological resistances of mosquitoes were quite stable across the villages and seasons within the spatiotemporal frame of the study. This contrasts with their biting rates, which was found, in another study (Taconet, Porciani, et al., 2021), highly variable across the villages, seasons, and amongst the species. This calls for distinct spatio-temporal management of interventions targeted at reducing human-vector contact and reducing resistance selection (both essential) in the field. While the former should be highly locally tailored (i.e. specific to each village and season) (Taconet, Porciani, et al., 2021), the latter, due to its stability across villages and seasons, would probably not benefit significantly from being customized at these spatio-temporal scales in our areas. In other words, while resistance management plans are undoubtedly urgently needed, there is no compelling evidence – in the current state of the knowledge - that they should be tailored at very fine scales (village, season). Noteworthy, mosquitoes were collected during the dry season and at the beginning and end of the rainy season, but, for logistical reasons, not at the peak of the rainy season (and therefore not at the likely peak of mosquito abundance). It would be worth collecting mosquitoes at this season to confirm the observed resistance rates.

## Conclusion

In an attempt to better understand the drivers of the intensity and spatio-temporal heterogeneity of physiological (genotypes) and behavioural (phenotypes) resistance in malaria vectors, at the scale of a rural health district over a period of 1.5 years, we have mainly (i) shown that resistance (both physiological and behavioural) was quite homogeneous across the villages and seasons at theses scales, and (ii) hypothesized that at these spatiotemporal scales, vector resistance seemed to be only marginally driven by environmental factors other than those linked to insecticide use in current vector control. Following the distribution of LLINs, the rapid widespread of physiological resistance occurring in tandem with probable lower acting behavioural adaptations, are very likely contributing to the erosion of insecticide efficacy on malaria vectors. We believe that without waiting to understand precisely the underlying drivers, mechanisms, and rates of selection of resistances, the malaria control community needs to think very strategically about the use and usefulness of current and novel insecticide-based control interventions.

## Acknowledgements

We thank populations of the villages for their kind support and collaboration. We also thank all the field and laboratory staff for their strong commitment to the REACT project.

## Abbreviations

AIC: Akaike Information Criterion
AUC: Area under the ROC Curve
BF: Burkina Faso
CV: Cross-Validation
GLMM: Generalized Linear binomial Mixed-effect Model
GPM: Global Precipitation Measurement
HLC: Human Landing Catch
IC: Ivory Coast
IRS: Indoor Residual Spraying
LLIN: Long Lasting Insecticide Nets
ML: Machine Learning
MODIS: Moderate Resolution Imaging Spectroradiometer
RF: Random Forest
SD: Standard Devia-tion
SR: Spatial Resolution
SSD: Spatial Standard Deviation
TSD: Temporal Standard Deviation
TR: Temporal Resolution
VC: Vector Control

## Data, scripts, code and supplementary information availability

Data and scripts are available online: https://doi.org/10.23708/LV8GEW (Taconet, D Soma, et al., 2023a) Supplementary information are available online: https://doi.org/10.23708/VJEEMU (Taconet, D Soma, et al., 2023b)

## Ethics approval and consent to participate

Ethical clearance for the study was granted by the National ethics committee (No. 063/MSHP/CNER-kp) in Côte d*’*Ivoire and by the Institutional Ethics Committee of the Institut de Recherche en Sciences de la Santé (No. A06/2016/CEIRES) in Bukina Faso. We received community agreement before the beginning of the study, and we obtained written informed consent from all the mosquito collectors and supervisors. Yellow fever vaccines were administered to all the field staff. Collectors were treated free of charge when they were diagnosed with malaria during the study period according to WHO recommendations. They were also free to withdraw from the study at any time without any consequences.

